# Uncovering Photolyase/Cryptochrome Genes Diversity in Aquatic Microbiomes Exposed to Diverse UV-B Regimes

**DOI:** 10.1101/701565

**Authors:** Daniel G. Alonso-Reyes, Maria Eugenia Farias, Virginia Helena Albarracín

**Author notes:** Corresponding author: Virginia Helena Albarracín, Centro Integral de Microscopía Electrónica (CIME, CONICET, UNT) Camino de Sirga s/n. Finca El Manantial, Yerba Buena (4107). Tucumán, Argentina.

## Abstract

During evolution, microorganisms exposed to high UV-B doses developed a fine-tuned photo-enzymes called “photolyases” to cope with DNA damage by UV-B. These photoreceptors belonging to the Cryptochrome/Photolyase Family (CPF) were well characterized at the genomic and proteomic level in bacteria isolated from a wide range of environments. In this work, we go further towards studying the abundance of CPF on aquatic microbial communities from different geographic regions across the globe. Metagenomics data combined with geo-referenced solar irradiation measurements indicated that the higher the UV-B dose suffered by the microbiome’s environment, the higher the abundance of CPF genes and lower the microbial diversity. A connection between CPF abundance and radiation intensity/photoperiod was reported. Likewise, cryptochrome-like genes were found abundant in most exposed microbiomes, indicating a complementary role to standard photolyases. Also, we observed that CPFs are more likely present in dominant taxa of the highly irradiated microbiomes, suggesting an evolutionary force for survival and dominance under extreme solar exposure. Finally, this work reported three novel CPF clades not identified so far, proving the potential of global metagenomic analyses in detecting novel proteins.

## 1. INTRODUCTION

Solar radiation is an essential factor sustaining complex life on Earth. In the past decades, the drastic reduction of stratospheric ozone produced by increased concentrations of chlorofluorocarbons (CFCs) (Aucamp 2007) and other halogen gases in the upper atmosphere (Russell III et al. 1996) resulted in a rise of biologically harmful UV radiation (UV) at the Earth’s surface together with its detrimental effect on all life forms.

The amount of UV radiation reaching the ground comprises only a small proportion of global radiation, about 6–7 % of UV-A (320–400 nm), less than 1.0 % of UV-B (280–315 nm) and virtually no UV-C (Hu et al. 2008). Biological damage is wavelength-dependent: UV-A produces mainly reactive oxygen intermediates causing indirect damage to DNA, proteins, and lipids. On the other hand, UV-B and UV-C (100 to 280 nm) cause both indirect and direct damage to DNA because of its strong absorption at wavelengths below 320 nm (Mitchell & Karentz 1993). In accordance, numerous studies reported the UV-B effects on plants (Searles et al. 2001, Xiong & Day 2001, Robinson et al. 2005, Ruhland et al. 2005, Jansen et al. 2010, Yan et al. 2012), animals (Robson et al. 2005, Bao et al. 2014), microorganisms (Zaller et al. 2002, Avery et al. 2004, Rinnan et al. 2005, Piccini et al. 2009) and ecosystems (Garcia-Pichel 1994, Karentz 1995, Joux et al. 1999, Ballaré et al. 2011, Häder et al. 2011).

Solar UV radiation can damage aquatic organisms and decrease the productivity of aquatic ecosystems. Beyond the targets for damage by UV radiation (DNA, lipids, protein) that are common for all biological systems, a major site of damage in primary producers is the photosynthetic machinery including photosystem II and the accessory pigments that funnel light energy to the reaction centers (Häder & Gao 2015). The subsequent damage will directly reduce primary production. Also, the effects of UV radiation may reduce the photosynthetic uptake of atmospheric carbon dioxide and affect species diversity, ecosystem stability, trophic interactions, and global biogeochemical cycles (Häder et al. 2011, Williamson et al. 2019).

During evolution, microorganisms exposed to high UV doses and other DNA damaging factors have developed specific and highly conserved DNA repair mechanisms. The most important are photoreactivation, excision repair (NER and BER), mismatch repair (MMR), and homologous repair (HR). Also, damage tolerance (dimer bypass), SOS response, checkpoint activation, and programmed cell death or apoptosis efficiently act against DNA lesions ensuring genomic integrity.

The finest and efficient mechanism to repair DNA damage by UV-B is photoreactivation, executed by photoreactivating enzymes known as “photolyases,” which target the main products of UV-B, cyclobutane pyrimidine dimers (CPD). Such CPD lesions bring polymerases to a standstill, eventually leading to cell death. These enzymes bind tightly to CPDs in the dark and can be activated by different wavelengths, causing the split of the two C–C bonds in the CPD unit and resulting in the re-formation of the two separate pyrimidine bases (S Weber 2005). Photolyases, together with the related cryptochromes (Cry), constitute the Cryptochrome/Photolyase Family (CPF). However, cryptochromes have no photolyase activity and function as signaling molecules regulating diverse biological responses such as entrainment of circadian rhythms in plants and animals (Roenneberg & Merrow 2005, Harmer 2009). It has been argued that CPF had had an early evolution in the history of life due to the need of the primitive organisms to cope with the exacerbated solar radiation in the time when no stratospheric ozone was protecting the Earth (Toh et al. 1997, Portero et al. 2019).

Investigations at metagenome-level about the effects of sunlight on microbial communities were mostly confined to photoreceptors such as microbial rhodopsins (Pushkarev & Béjà 2016, Pushkarev et al. 2018) and LOV-domains (Pathak et al. 2012). Likewise, Singh et al. (2009) reported abundances for several light-related genes in microbiomes from different environments (Singh et al. 2009). However, abundance and diversity of CPF on microbiomes in a world-wide scale and light range were not studied so far, although being these photoreceptors a vital issue in the defense against UV-B. In this work, we studied aquatic microbiomes exposed to different intensities of UV-B and photoperiods around the globe through metagenomic analysis of existing published DNA datasets. Thus, the objective was to characterize the occurrence and diversity of CPF on microbiomes from aquatic ecological niches, with some of them suffering from extreme conditions. Our results add a new dimension to the understanding of the short-term effects of climate change and atmospheric ozone depletion on environmental microbial communities. Besides, the focus of this work on microbial communities from extreme environments provides models for early life research.

## 2. MATERIALS AND METHODS

### 2.1 Metagenome selection and UV-B data processing

The datasets of metagenomes selected for this work from the NCBI database were obtained and sequenced by shotgun strategy through Illumina technology except for Lake Diamante metagenome, which was sequenced with 454 GS FLX Titanium instrument. The accession numbers are the following: Lake Diamante (DM) (ERR1824222), Socompa stromatolite (SS) (SRR3341855), Tibetan Plateau sediment (TB) (SRR3322106), Amazon River (AM) (SRR1790676), Lake Rauer (RA) (SRR6129205), Dewar Creek hot spring (HT) (SRR5580900), Greenland cryoconite (CR) (SRR5275901), Lake Montjoie (MT) (SRR5818193), Olkiluoto Island groundwater (OK) (SRR6976411) and human gut (GU) (SRR6517782).

An essential aspect of this work was to reanalyze published metagenomic data of world-wide distributed microbiomes taking into account their overall UV-B exposure (Fig. 1). Monthly mean UV-B glUV datasets (Beckmann et al. 2014) processed using QGIS (www.qgis.org) were utilized to calculate UV-B exposure of the samples of the selected metagenomes according to the date/location of sampling available at the “biosample” linked section of each SRA entry. To compensate for the lack of replicates on most samples, we performed the analysis by clustering the datasets in three groups according to UV-exposure regimes: High (UV_High_), Mid (UV_Mid_), and low-exposed (UV_Low_). The metagenome of the human gut was assumed as a UV-B free environment, acting as a negative control. Table 1 showed the main characteristics of these environmental datasets when available. A brief description of the environmental conditions of each sampling site is summarized below, including citation to previous works describing the corresponding microbial communities with further detail:

**Figure 1.**
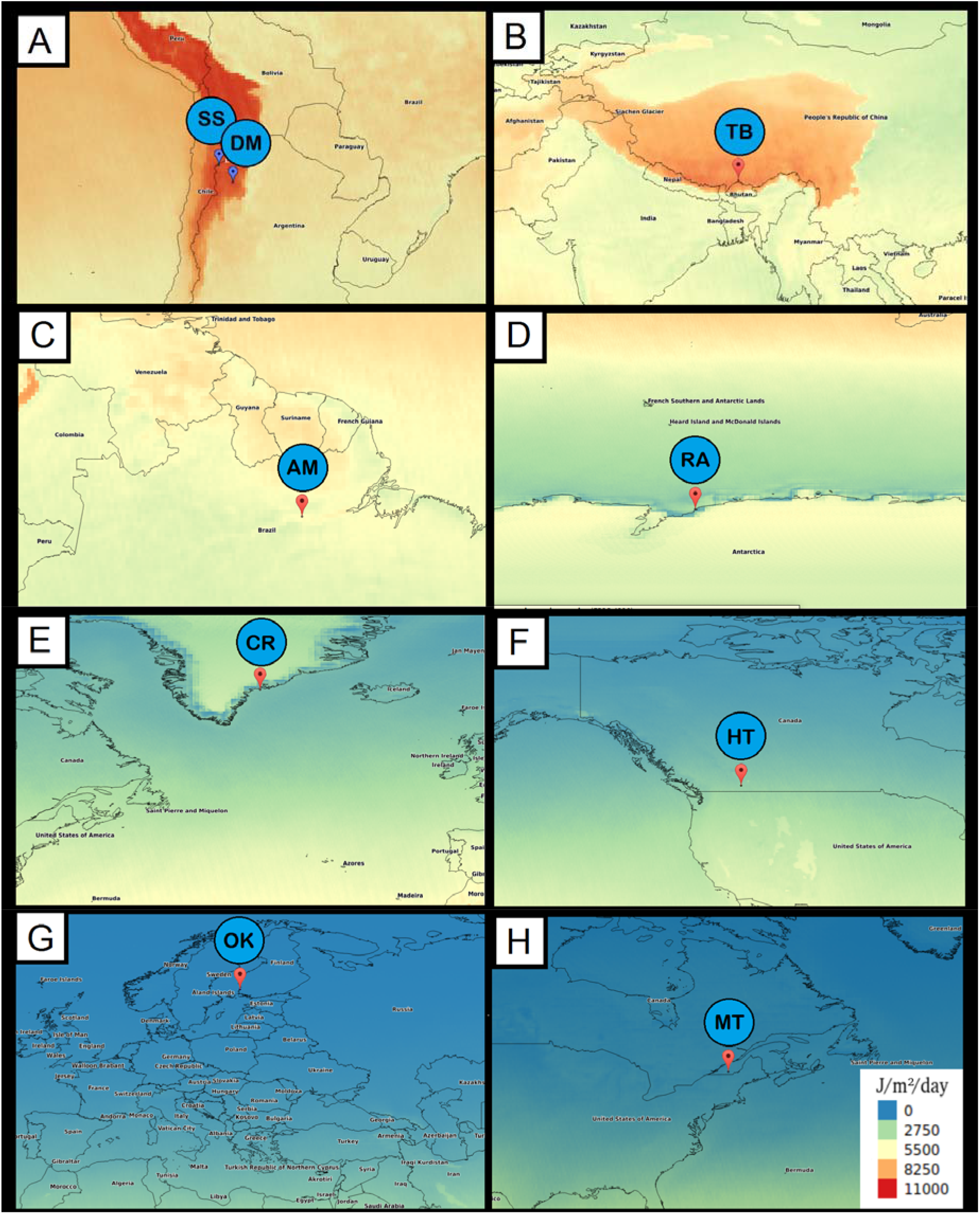
Worldwide metagenomic sampling linked to UV-B georeferenced data obtained from glUV datasets (https://www.ufz.de/gluv/). The metagenomes were selected from NCBI using biosample information: DM, Lake Diamante; SS, Socompa stromatolite; TB, Tibetan Plateau sediment; AM, Amazon River; RA, Lake Rauer; HT, Dewar Creek hot spring; CR, Greenland cryoconite; MT, Lake Montjoie; OK, Olkiluoto Island groundwater.

**Table 1.**
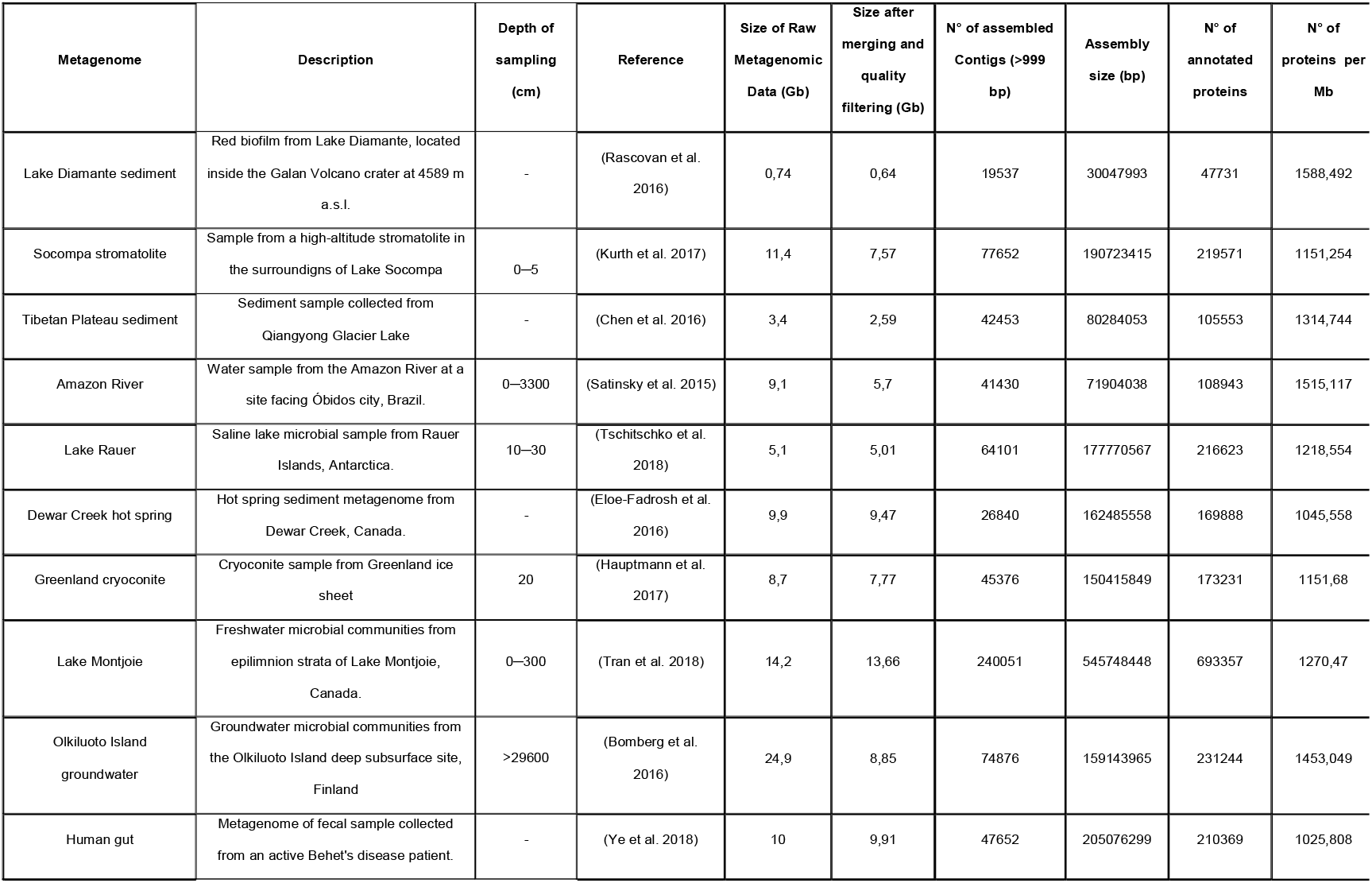
General traits of the metagenomes used in the current study.

| Metagenome | Description | Depth of sampling (cm) | Reference | Size of Raw Metagenomic Data (Gb) | Size after merging and quality filtering (Gb) | N° of assembled Contigs (>999 bp) | Assembly size (bp) | N° of annotated proteins | N° of proteins per Mb |
| --- | --- | --- | --- | --- | --- | --- | --- | --- | --- |
| Lake Diamante sediment | Red biofilm from Lake Diamante, located inside the Galan Volcano crater at 4589 m a.s.l. | - | (Rascovan et al. 2016) | 0,74 | 0,64 | 19537 | 30047993 | 47731 | 1588,492 |
| Socompa stromatolite | Sample from a high-altitude stromatolite in the surroundings of Lake Socompa | 0–5 | (Kurth et al. 2017) | 11,4 | 7,57 | 77652 | 190723415 | 219571 | 1151,254 |
| Tibetan Plateau sediment | Sediment sample collected from Qiangyong Glacier Lake | - | (Chen et al. 2016) | 3,4 | 2,59 | 42453 | 80284053 | 105553 | 1314,744 |
| Amazon River | Water sample from the Amazon River at a site facing Óbidos city, Brazil. | 0–3300 | (Satinsky et al. 2015) | 9,1 | 5,7 | 41430 | 71904038 | 108943 | 1515,117 |
| Lake Rauer | Saline lake microbial sample from Rauer Islands, Antarctica. | 10–30 | (Tschitschko et al. 2018) | 5,1 | 5,01 | 64101 | 177770567 | 216623 | 1218,554 |
| Dewar Creek hot spring | Hot spring sediment metagenome from Dewar Creek, Canada. | - | (Eloe-Fadrosh et al. 2016) | 9,9 | 9,47 | 26840 | 162485558 | 169888 | 1045,558 |
| Greenland cryoconite | Cryoconite sample from Greenland ice sheet | 20 | (Hauptmann et al. 2017) | 8,7 | 7,77 | 45376 | 150415849 | 173231 | 1151,68 |
| Lake Montjoie | Freshwater microbial communities from epilimnion strata of Lake Montjoie, Canada. | 0–300 | (Tran et al. 2018) | 14,2 | 13,66 | 240051 | 545748448 | 693357 | 1270,47 |
| Oikiluoto Island groundwater | Groundwater microbial communities from the Oikiluoto Island deep subsurface site, Finland | >29600 | (Bomberg et al. 2016) | 24,9 | 8,85 | 74876 | 159143965 | 231244 | 1453,049 |
| Human gut | Metagenome of fecal sample collected from an active Behet's disease patient. | - | (Ye et al. 2018) | 10 | 9,91 | 47652 | 205076299 | 210369 | 1025,808 |

DM samples were taken from red biofilms attached to the bottom of calcareous stones submerged in Lake Diamante on 15^th^ February 2012. This lake is located inside the Galan Volcano crater placed at 4,589 m above sea level (coordinates: −26.008889, - 67.043333). High pH (9 to 11), high arsenic concentrations (115 to 234 mg 1 – 1), high salinity (270 g 1 − 1, 217 mS cm – 1), high UV radiation (84 W m −2 of UVA-B at noon), long day-night temperature range (−20 °C to +20 °C) and low O_2_ pressure constitutes a set of unique extreme conditions that prevails in the lake (Rascovan et al. 2016).

SS belongs to stromatolites found along the southern shore of Socompa Lake (-24.5361333, −68.20275) at the base of the Socompa Volcano (3,570 masl). These structures were found around the border of Socompa Lake during the summer when they are partially exposed depending on the tide and hydrological regime; at the opposite, they are entirely submerged during winter and spring. The stromatolites are rounded domal structures that present a clear stratification that appear in vertical sections. The samples were cylinders of 5 cm deep from the top of the stromatolite. The site was exposed to air at the moment of sampling in February 2011. Extreme environmental conditions of the lake include hypersalinity, high thermal amplitude with daily temperatures that range from −10 °C to 20 °C in summer and −20 °C to 10 °C in winter, UV solar irradiance that reaches 68 W m-2 19, 32, low O_2_ pressure, low nutrient availability and, primarily, high arsenic content (18.5 mg L-1) (Kurth et al. 2017).

TB was collected on 1^st^ August 2013 from sediments of the Qiangyong Glacier Lake (28.51, 90.12) that is located in the southern part of the Tibetan Plateau, in the Indian monsoon climate region. The temperature, pH, and electric conductivity of surface water are, on average, 6.89 °C, 8.32, and 136.4 μS cm^-1^, respectively. (Chen et al. 2016).

AM water sample was collected on 19^th^ May 2011 from Amazon River at Obidos station (−1.919017, 55.525717) at 33 m of depth by pumping water with a Shurflo submersible pump through a 297 μm stainless steel screen. The temperature, pH, and O_2_ were 28.4 °C, 6.6, and 3.2 m/L respectively on the site of sampling (Satinsky et al. 2015, Doherty et al. 2017).

RA was sampled on 12 January 2015 from Rauer Lake (Torckler Island; −68.5558 78.1913). Water temperature and approximate depth of water at the sampling site were 9°C and 10-15cm. The lake was shallow <30cm. There was a crust of salt crystals on the sediment (which make the lake look frozen from the air). There also appeared to be stratification in the lake: a top clear layer of ~10cm and a bottom layer of ~5cm. These layers weren’t visible until they were disturbed, and a visible haze produced when these layers were mixed. (Tschitschko et al. 2018)

CR belongs to a cryoconite sample collected on 29^th^ August 2013 from Greenland ice sheet in the TAS-U-A1 site (65.41, −38.51). pH, altitude, and depth of sampling were 5.53, 580 m, and 20 cm, respectively. Cryoconite was collected from 2–18 holes at each site, depending on their availability, using sterile 50 ml syringes (Stibal et al. 2015a b, Hauptmann et al. 2017).

MT microbial sample was collected from Lake Montjoie, in Canada (45.4091, - 72.0994 W) on February 2014, being the lake covered of ice at the time of sampling. The sample was taken from the epilimnion strata of the lake (0-3 m), and pH in the site of sampling was 6.41 (Tran et al. 2018).

HT sediment sample was collected from Dewar Creek hot spring (49.9543667, - 116.5155000) near the source of the hot spring on 28^th^ September 2012. pH and temperature of the environment were 7.93 and 66.4 °C (Eloe-Fadrosh et al. 2016).

OK belongs to groundwater collected from a deep subsurface site KR11_0.1 (61.2413, 21.4947) in Olkiluoto Island, on the south-west coast of Finland on 9^th^ September 2016. Given the environmental data provided by the authors of the original research (Bomberg et al. 2016), 60 drill holes were collected from different fractures at different depths (296-798 mbsl). The groundwater is stratified with a salinity gradient extending from fresh to brackish water to a depth of 30 m and the highest salinity concentration of 125 g L-1 total dissolved solids (TDS) at 1,000 m depth (Posiva, 2013). Between 100 and 300 m depths, the groundwater originates from ancient (pre-Baltic) seawater and has high concentrations of SO_2_^-4^. The temperature rises linearly with depth, from ca. 5–6 °C at 50 m to ca. 20 °C at 1,000 m depth (Ahokas et al., 2008). The pH of the groundwater is slightly alkaline throughout the depth profile. The bedrock of Olkiluoto consists mainly of mica gneiss and pegmatitic granite-type rocks (Kärki and Paulamäki, 2006). The *in-situ* temperature at 300 m depth in the Olkiluoto bedrock is stable at approximately 10 °C and increases linearly to approximately 16 °C at 800 m depth.

GU corresponds to a gut metagenome collected from the faeces of an active Behcet’s disease patient (Ye et al. 2018) was used as negative control with no exposure to UV.

### 2.2. Meta-analysis, quality control, and assembly of selected metagenomes

Quality filtering and merging yielded a range between 0.64 and 13.66 Gb for further analysis (Table 1). Only TB and AM datasets reported a high percentage of merging with FLASH (Magoč & Salzberg 2011); however, the low-merged dataset from OK was also used for downstream steps as it ended with a reasonable size (between the range mentioned above). MEGAHIT (Liu et al. 2015) was used for assembly since it can deal with large and complex datasets in a time- and cost-efficient manner. Protein prediction with Prodigal (Hyatt et al. 2010) over contigs with >999 bp outputted a rate of 1000-1500 proteins / Mb, which is congruent with the high coding density expected for microbial species.

Adapters were removed from Illumina raw reads using fastq-mcf tool of ea-utils v1.04.676 (Aronesty 2011). This step was not needed for Lake Diamante metagenome, as it was sequenced with 454 technologies. The quality filtering and trimming were performed with the same program using parameters l>50 and q>20. The program kneaddata v0.6.1 (https://bitbucket.org/biobakery/kneaddata/wiki/Home) was used with the --bypass-trim option to clean human contaminant sequences. Pair end reads were merged with FLASH v1.2.11 (Magoč & Salzberg 2011) to recover unpaired longer reads. The pair end reads with a low percentage of merging were left in the paired state for assembly. Assembly of filtered reads was performed with MEGAHIT v1.1.2 (Liu et al. 2015). Assembled contigs were annotated with Prodigal v2.6.2 (Hyatt et al. 2010), which outputted translated protein sequences.

To perform quantitative metagenomics, pair-end reads that remained with a low percentage of merging were concatenated together by the forward and reverse using a script available at GitHub (https://github.com/LangilleLab/microbiome_helper/blob/master/concat_paired_end.pl).

### 2.3. Reference database building

Considering the main functional orthologous of the CPF group, a reference protein database was built. The two KEGG Orthology number corresponding to CPF members (K06876, K01669) was linked with the UniProt database, and then filtered results by EC number, known molecular function or biological process, uniref 90 clusters, and sequence length >100 subsequently. The single, concatenated, and merged reads were aligned to the aforementioned databases with PALADIN, which outputted read counts. The counts were normalized to Reads per Kilobase per Genome, and the number of genomes computed through MicrobeCensus v1.1.0 (Nayfach & Pollard 2015). The following formula describes the normalization:

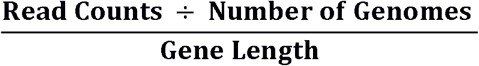

### 2.4. Estimation of gene abundance

To search and quantify the abundance of the genes in the different metagenomes, we aligned the preprocessed unpaired reads against the CPF reference database created before. The protein alignment was performed with PALADIN v1.3.1 (Westbrook et al. 2017), which outputs a table with alignment counts for each gene. Alignments were filtered by maximum quality = 60.

Diversity analysis of each metagenome was performed with MetaPhlAn v2.7.7 (Truong et al. 2015) with the --bt2_ps parameter set to ‘sensitive.’ The program MetaPhlAn profiles the composition of microbial communities from metagenomic shotgun sequencing data at species-level. It relies on ~1M unique clade-specific marker genes identified from ~17,000 reference genomes (~13,500 bacterial and archaeal, ~3,500 viral, and ~110 eukaryotic). Shannon diversity index was obtained with Vegan 2.5-2 R package (Oksanen et al. 2018).

### 2.5. Analysis of the annotated genes

The protein sequences were aligned using Diamond v0.9.22 software (Buchfink et al. 2014) against the CPF genes of the reference database built earlier. Alignment parameters were 50% identity, 70% query coverage and e < 10^-5^. The retrieved files coming from each metagenome were modified at the sequence headers to hold the name of its respective metagenome. Next, all files were merged in a single one and filtered by sequence length > 400 residues.

The filtered sequences were used for phylogenetic analysis. The phylogenetic tree was built with FastME v2.0 (Lefort et al. 2015) using the Jones–Taylor–Thornton rate matrix (Jones et al. 1992) with 1000 bootstrap replicates. A consensus of the 1000 resulting trees was selected for further processing and visualization using iTOL (Letunic & Bork 2016). Additionally, a taxonomical identification of the sequences was performed through BLAST web server (https://blast.ncbi.nlm.nih.gov/Blast.cgi).

## 3. RESULTS AND DISCUSSION

### 3.1. UV-B intensity profiles on worldwide microbiomes

The UV-B geo-referenced data helps us to group microbiomes according to their UV-exposure regimes in UV_High_, UV_Mid_, and UV_Low_ as described in materials and methods. The UV_High_ included DM, SS, and TB with UV-B intensities of 9677, 9536, and 8885 J/m2/day, respectively. In contrast, UV_Low_ microbiomes OK, MT, and CR suffered from low UV-B intensities of 77, 719, and 1759 J/m2/day, respectively. In between the UV_Mid_ microbiomes AM, RA, and HT, received intensity of 5630, 2289, and 2281 J/m2/day, respectively. Important to remark is that these intensity values were registered taking in account the sampling dates; for instance, the samples of the UV_High_ group were registered during summer, when the insolation is maximum on the high altitude environments of the Argentinean Puna and the Tibetan Plateau (February and August), and UV-B incidence become the highest of the planet. On the other hand, the UV_Low_ microbiomes were sampled during August, November, and February, which correspond to summer, autumn and winter seasons in the northern hemisphere with lower levels of irradiation as compared to the southern hemisphere. In turn, the UV_Mid_ included the AM microbiome from a tropical environment, and RA and HT from high-latitudes extreme cold and hot environments, respectively.

The calculated day-length from the UV-datasets for each georeferenced metagenome (https://www.suncalc.org) revealed that two microbiomes suffered from peculiar photoperiods at the time of sampling. While the OK microbiome was experiencing a short photoperiod-only 7.8 hours-, RA was sampled when Antarctica went through an entire day under the sun. We assumed that the environments were exposed to similar conditions of indicated solar irradiance for a considerable broad period already before sampling. In consequence, the diversity of species and genes on genomes are a reflection of the ecological pressure of radiation on the environment (among other factors) as the geographic conditions (latitude, altitude, and orography) did not change considerably in those sampled regions for decades. Although no available, transcriptomic data on these sites would be an excellent tool to evaluate the reflection of environmental pressure on the dynamics of the microbial communities. Besides, the intensity of solar irradiation over each microbiome also depend on their on-site spatial disposition being maximum for samples taken from soils or shallow water of lakes, springs, oceans, and streams but much lower in sediments, groundwater or deep water.

It is known that UV-B negatively affects microbial diversity (Ballaré et al. 2011); nevertheless, this was not assessed so far in a systematic way across different irradiation regimes on a global scale. The herein results used disperse metagenomic data to assessed species richness and Shannon diversity index in diverse ecological niches, suggest an interesting trend (Fig. 2); TB, SS, and DM from the UV_High_ group had the lowest species number, while the GU microbiome, which received null irradiation, had the maximum number of species. Moreover, microbiomes exposed to intensities below 4000 J/m^2^/day, revealed relatively high species richness (number of species: 11-103), while SS microbiome with intensities above 8000 J/m2/day only showed 21 species.

**Figure 2.**
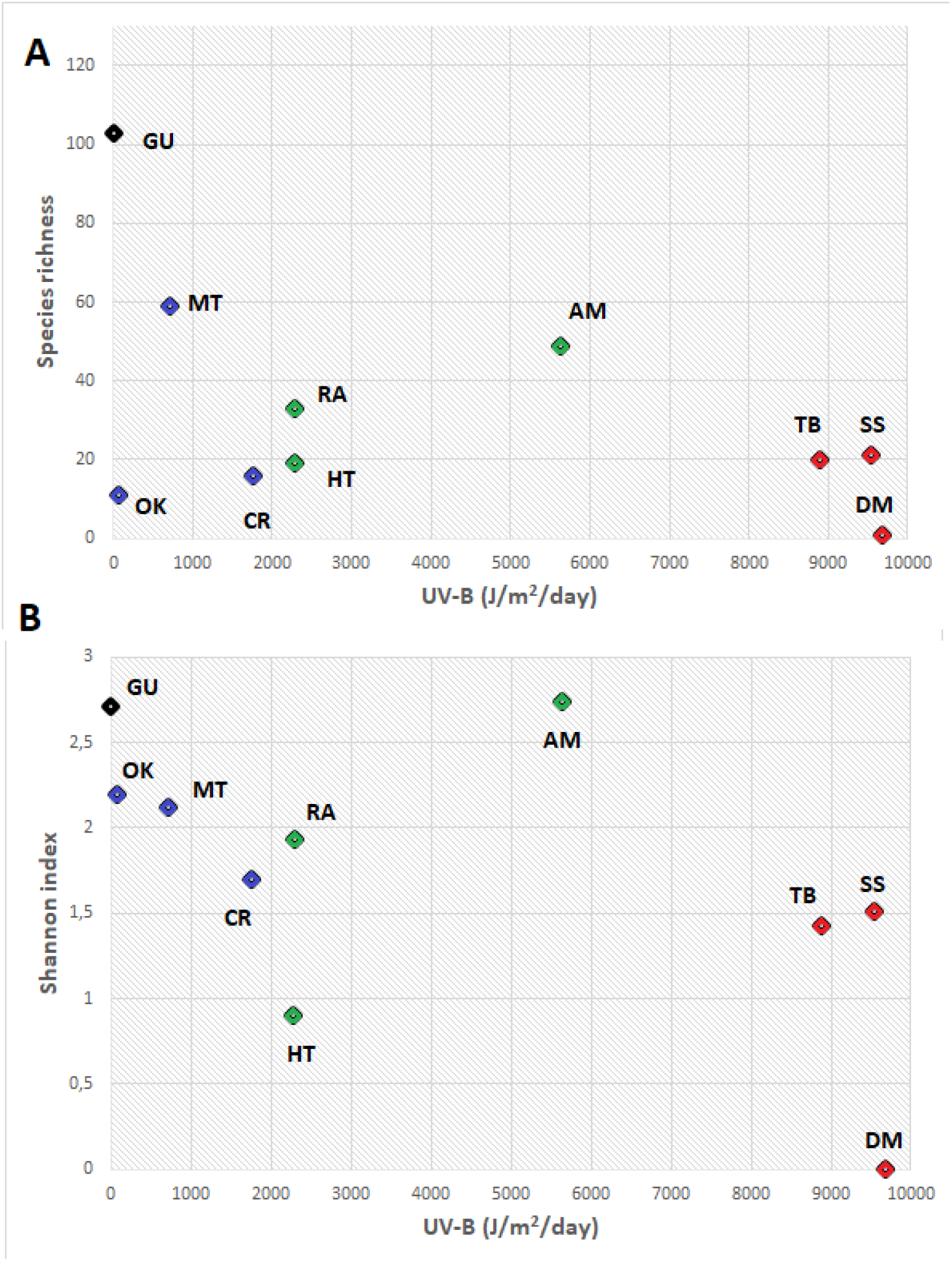
Scatterplots showing the impact of UV-B on the microbial diversity in the whole metagenomic dataset. Two parameters were measured: species richness (A) and the Shannon index (B). Metagenomes were also colored according the group they belong, being red for UV_High_, green for UV_Mid_ and blue for UV_Low_. Abbreviations: DM, Lake Diamante; SS, Socompa stromatolite; TB, Tibetan Plateau sediment; AM, Amazon River; RA, Lake Rauer; HT, Dewar Creek hot spring; CR, Greenland cryoconite; MT, Lake Montjoie; OK, Olkiluoto Island groundwater; GU, human gut.

Species richness (Fig. 2A) was quite variable for microbiomes whose UV-B exposure was below a certain threshold (6000 J/m^2^/day). In microbiomes receiving higher UV doses, the number of species decreased, indicating selective pressure. Those species that do not possess efficient molecular mechanisms to defend themselves from UV-B become unable to adapt to the new assemblage of species. A similar situation was observed when applying the Shannon index (Fig. 2B) (which incorporates equitability of the species abundance in addition to its ability to detect rare species): microbiomes of the UV_High_ group, and HT (UV_Mid_) presented the lowest Shannon index values. These results suggest that those microorganisms with full capacity to defend themselves against UV-B radiation reach dominance, relegating others less fitted. Nevertheless, this correlation is not conclusive as many other chemical and physical factors (salinity, nutrients availability, heavy metals) may be causing selective pressure on microbial diversity.

### 3.2. Quantitative analysis of CPF

In order to evaluate the abundance of CPF genes in each microbiome, a database was built using Uniprot sequences linked to KEGG orthology and then aligned to the query metagenomes. Our results show that the distribution of CPF genes in metagenomes followed a predicted ecological pattern; CPF genes were abundant in microbiomes with UV_High_ and UV_Mid_, while insignificant or null in UV_Low_ microbiomes-OK and GU respectively. In DM, CPF genes appeared profusely. Their abundances were 249 e^-02^ RPKG, which were higher than in the rest of the microbiomes.

The relationship between the abundance of CPF and intensity/time of insolation was studied (Fig. 3A), showing an upward trend of CPF abundance as UV-B irradiation increases. However, we have seen that RA, an UV_Mid_ microbiome, completely deviates from the trend having because of its high abundance. We had previously mentioned that RA has the most extended photoperiod of the study, so we set out to verify if there is a connection between the abundance of CPF and the photoperiod. Figure 3B shows that there is an upward trend of CPF as the photoperiod increases, finding RA quite in line with the trend. Thus, both factors, the intensity, and the photoperiod could be influencing the abundance of CPF in the microbial communities

**Figure 3.**
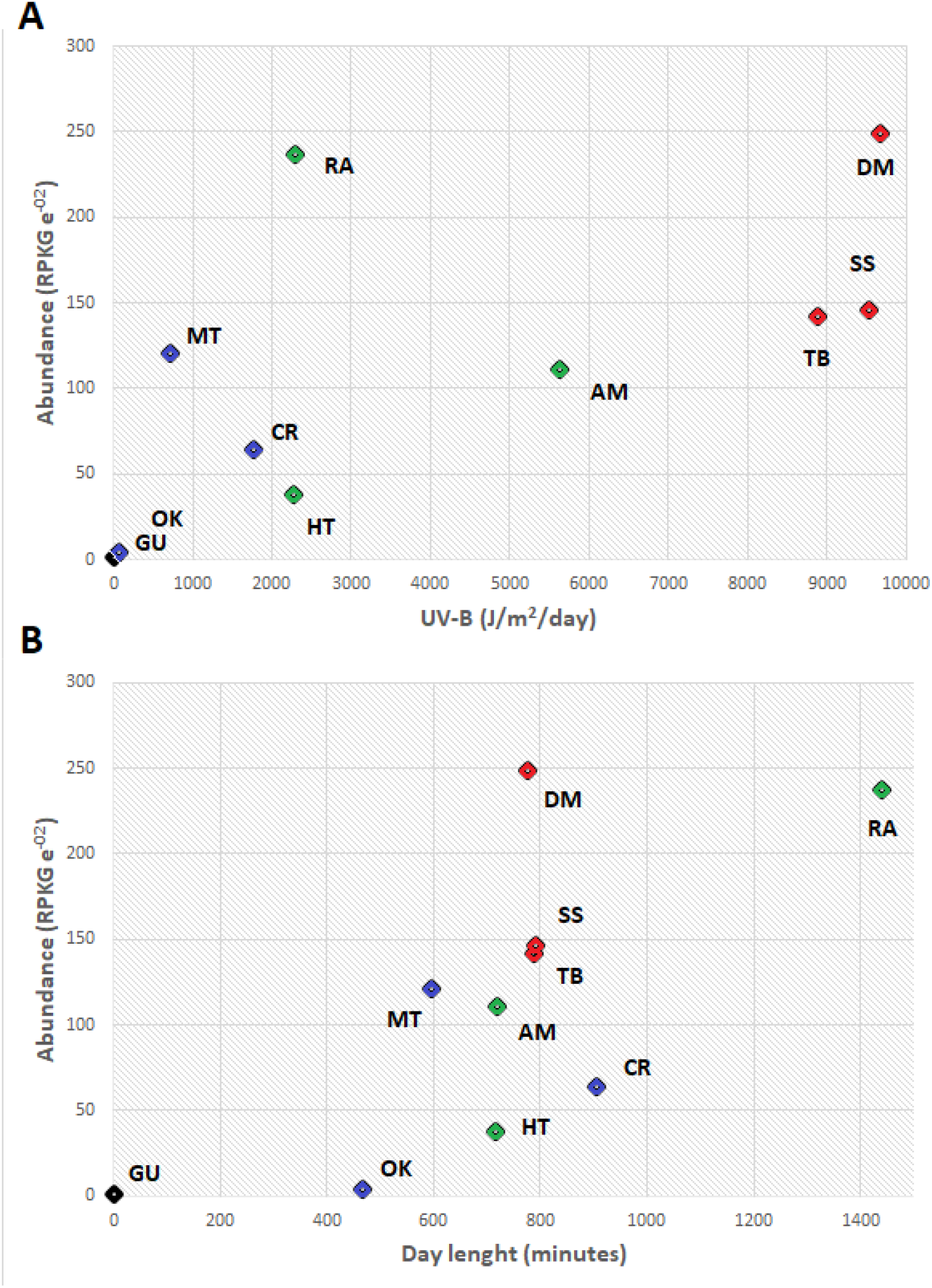
Scatterplots showing the impact of UV-B (A) and Day length (B) on the CPF abundance in the whole metagenomic dataset. Metagenomes were also colored according the group they belong, being red for UV_High_, green for UV_Mid_ and blue for UV_Low_. Abbreviations: DM, Lake Diamante; SS, Socompa stromatolite; TB, Tibetan Plateau sediment; AM, Amazon River; RA, Lake Rauer; HT, Dewar Creek hot spring; CR, Greenland cryoconite; MT, Lake Montjoie; OK, Olkiluoto Island groundwater; GU, human gut.

Our study reveals a rising trend of CPF abundance in microbial communities as their environments receive higher radiation or day duration extends. That is likely due to an increase in DNA damage caused by UV-B, which are mainly pyrimidine dimers. In such a situation, populations could increase the expression of CPF by increasing the gene copy number. Besides, species lacking CPF in their genome would be replaced by species that contain these genes. In either case, the overall increase in CPF copy abundance would indicate that the community improves its defense capabilities against UVB using the highly efficient mechanism of photoreactivation or modulating enzymatic mechanisms triggered by light sensing by cryptochromes.

### 3.4. Phylogenetic analysis of CPF genes

The CPF group of proteins comprises mostly genes of photolyases with different specializations and cryptochromes whose functions are mostly unknown. The family has been divided into different subgroups considering phylogeny, chromophore type, specialization, host organism, and structure. Alignment, editing, and subsequent phylogenetic analysis of 214 CPF sequences found in the metagenomes configured a robust tree with numerous clades (Fig. 4). The sequences mostly grouped within well-known subfamilies: the CPD photolyases classes I, II, and III, DASH cryptochromes or single-strand photolyases (Selby & Sancar 2006), the FeS-BCPs group that consist of proteins having an iron-sulfur cluster either prokaryotic [6-4] photolyases or cryptochromes (Graf et al. 2015) and the thermostable CPD photolyases, a group of unstable position in phylogeny, that have FMN as antenna cofactor and a thermostable nature (Ueda et al. 2005) (Portero et al. 2019). Also, this work reports three novel extra clades: i.e., unidentified I, II, and III (UI, UII, and UIII) with 41%, 88%, and 96% of bootstrap support, respectively. The three new clades grouped 24.76% of the whole sequences in the analysis (Fig. 5), thus offering us a large pool of candidates for new functions, specializations, or molecular specificities. Interestingly, the clades UI and UII are mainly composed of sequences of a single microbiome; sequences from the RA microbiome constitute 85.7% of UI, while 87.5% of UII corresponds to MT sequences. The UI subfamily is the only clade in DM but dominant in RA; both microbiomes displayed the highest content of CPF’s genes in this study. Furthermore, UI and UII clades are taxonomically homogeneous, with their sequences belonging to Halobacteria and Actinobacteria, respectively. On the other hand, UIII is formed by sequences of different taxonomic origins, including Proteobacteria, Bacteroidetes, and Verrucomicrobia phylum.

**Figure 4.**
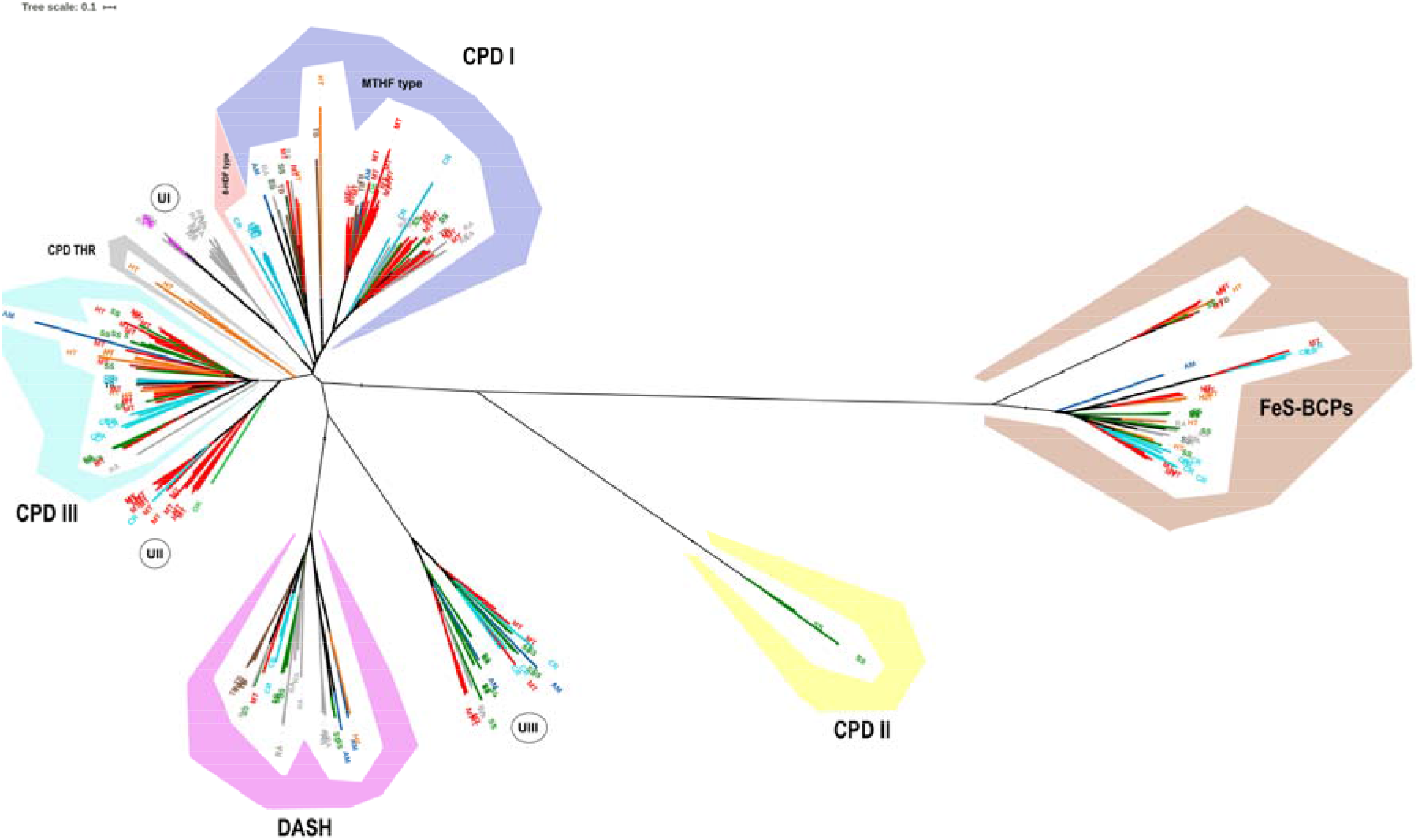
Phylogenetic tree built with the CPF sequences from the whole metagenomic dataset. Each leaf is colored and labeled by sample procedence: DM, Lake Diamante (purple); SS, Socompa stromatolite (dark green); TB, Tibetan Plateau sediment (brown); AM, Amazon River (blue); RA, Lake Rauer (grey); HT, Dewar Creek hot spring (orange); CR, Greenland cryoconite (cyan); MT, Lake Montjoie (red); OK, Olkiluoto Island groundwater (light green). Abbreviations: CPD I, II and III refers to class I, II and III cyclobutane pirimidine dimer photolyases; CPD THR, thermostable CPD photolyase; DASH, DASH cryptochrome; FeS-BCPs, FeS bacterial cryptochromes and photolyases; UI, II and III refers to unidentified group I, II and III respectively.

**Figure 5.**
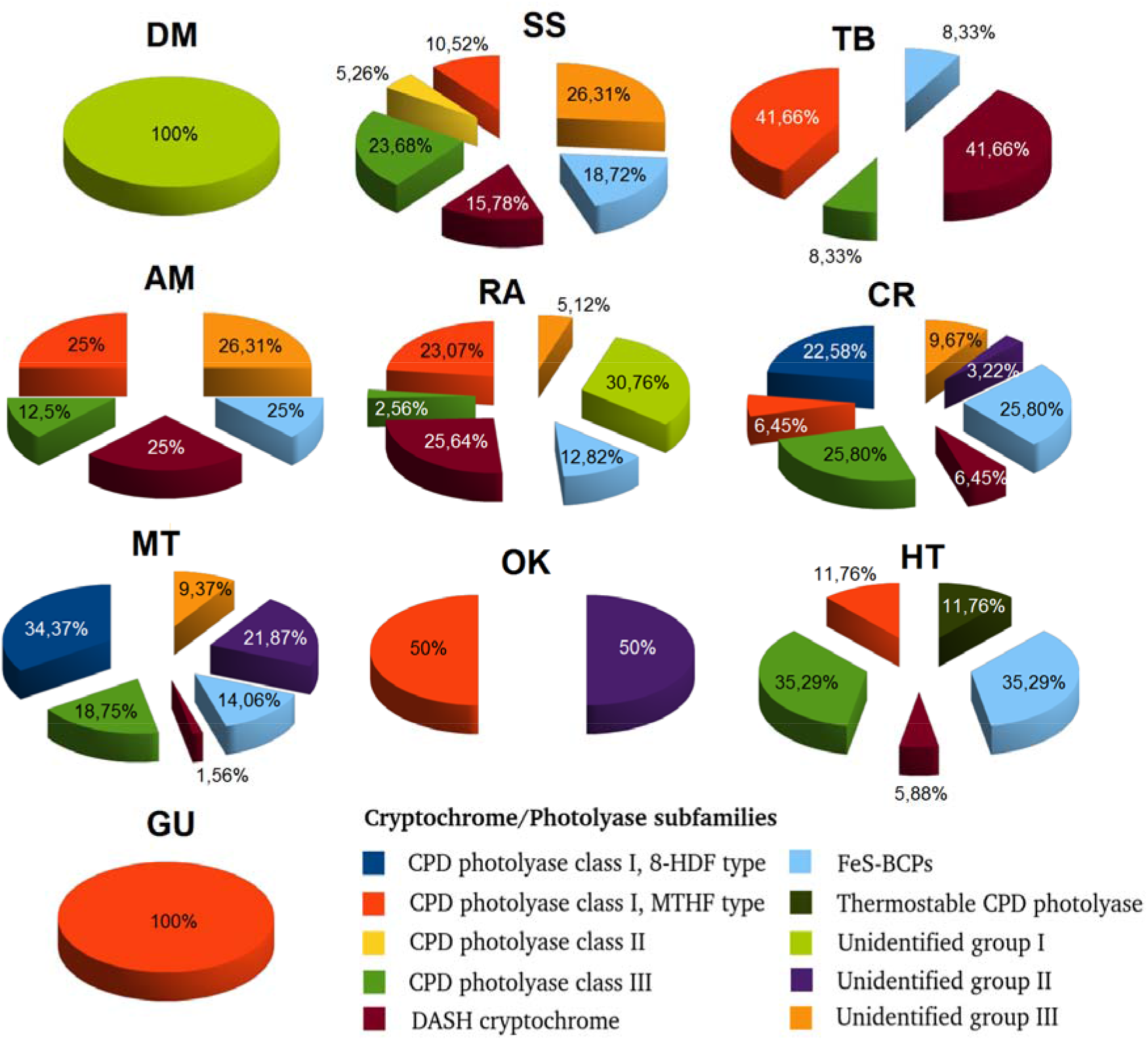
Distribution of each subfamily by metagenome. Abbreviations: DM, Lake Diamante; SS, Socompa stromatolite; TB, Tibetan Plateau sediment; AM, Amazon River; RA, Lake Rauer; HT, Dewar Creek hot spring; CR, Greenland cryoconite; MT, Lake Montjoie; OK, Olkiluoto Island groundwater; GU, human gut.

It is clear the widespread presence of cryptochrome-holding subfamilies cry-DASH and Fes-BCPs among the microbiomes (Fig. 6). The abundance of cry-DASH followed a pattern in consistence with higher irradiation conditions; the highest number of cry-DASH sequences are found in SS and TB communities (UVB_High_), AM (UVB_Mid_) and RA (the most prolonged photoperiod). It has been reported in previous works that these cryptochromes are, in fact, photolyases with an affinity for single-stranded DNA and, in some cases RNA (Selby & Sancar 2006). Cry-DASH may play a complementary role of standard photolyases with an affinity for double-stranded DNA, contributing to the global increase of photoreactivation in these microbiomes. The case for FeS-BCPs group is intriguing, being present in almost all the communities with similar relative abundance. It has been suggested that this class complements other photolyases by performing the function of a [6-4] photolyase (Zhang et al. 2013, Graf et al. 2015), avoiding or decreasing in this way the use of the inefficient NER system. Our work gives more support to such affirmation by showing its ubiquitous and homogenous presence in world-wide microbiomes.

**Figure 6.**
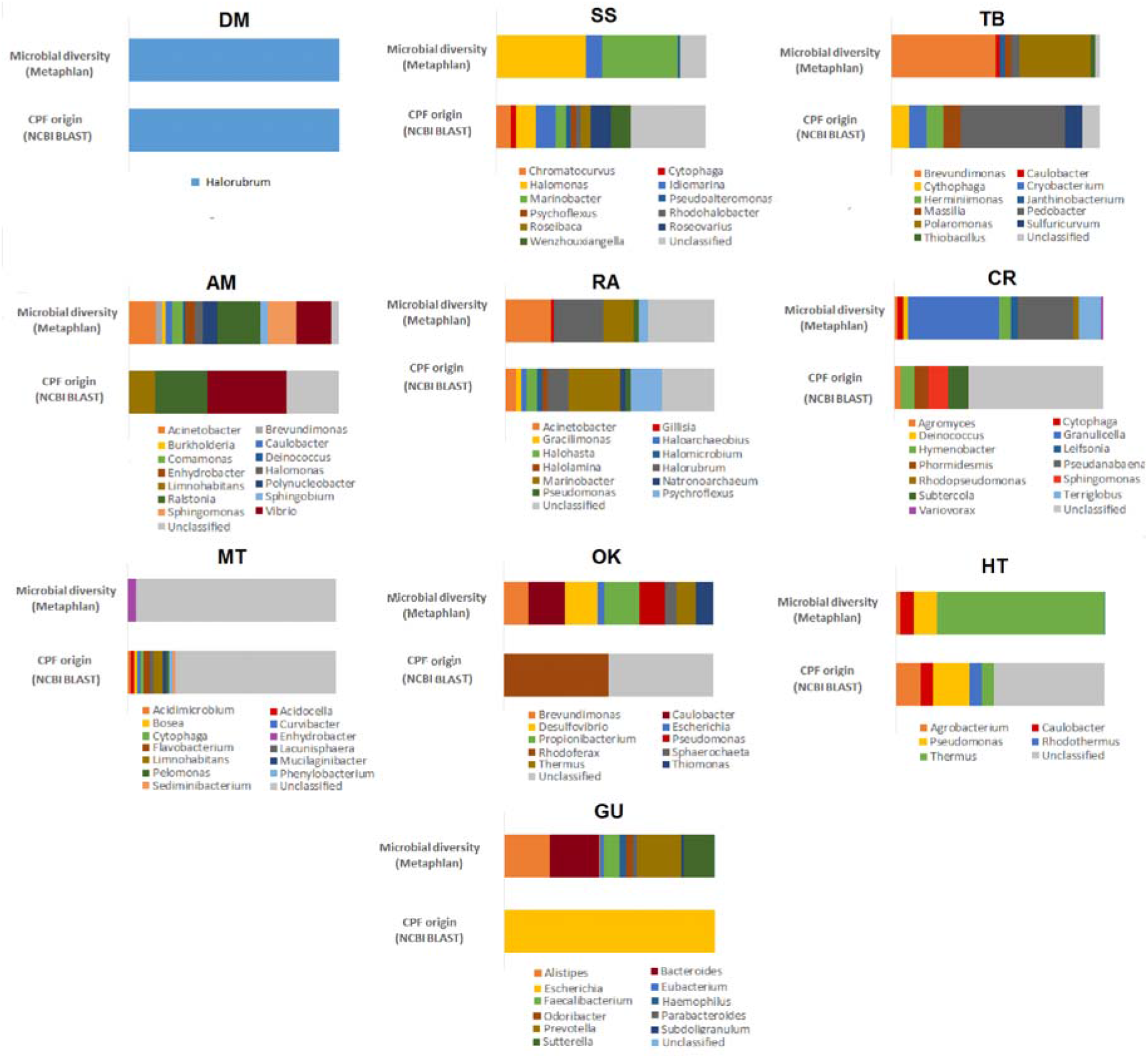
Comparative barplot showing for each metagenome the genus diversity both for whole microbial community (top bar) and the set of CPF sequences recovered from the same metagenome (lower bar). The diversity of the microbial community was estimated with Metaphlan, while the diversity of the set of CPF sequences were computed with Blast online platform. Abbreviations: DM, Lake Diamante; SS, Socompa stromatolite; TB, Tibetan Plateau sediment; AM, Amazon River; RA, Lake Rauer; HT, Dewar Creek hot spring; CR, Greenland cryoconite; MT, Lake Montjoie; OK, Olkiluoto Island groundwater; GU, human gut.

### 3.5. Relationship between CPF and taxa dominance

The assignment of CPF genes to a particular taxonomic group was also considered in this work. CPF sequences were classified by genus and paired with the same information previously obtained for each microbiome through MetaPhlAn. The CPF sequences grouped with their homolog with a BLAST identity lower than 70% clustered as an unclassified category. Genus diversity from the whole microbiome (top bar) and the set of CPF sequences recovered from that microbiome (lower bar) are shown (Fig. 6). The classification rate of the CPF genes was generally high (>60%) in DM, SS, TB, AM, RA, and GU but lower in the rest of the samples. MT had the lowest percentage of sequences (23%) assigned to some genus. This phenomenon may be a consequence of the poorly referenced diversity of the community itself (top bar). Despite this, MT had the most substantial diversity of genus (12) with CPF hits along with SS and RA (11 each).

The most abundant taxa in DM, SS, RA, and HT communities possess CPF genes. In AM, only *Ralstonia* and *Vibrio* displayed CPF; both genera together represented 35% of the community. In the case of CR, only *Hymenobacter* and *Agromyces* possessed these genes, with both groups adding up to 7%. *Pedobacter* was the unique taxon in TB with CPF genes, and it represented less than 4% of the community. Finally, *Escherichia*, with an abundance of just 0.44%, was the unique CPF contributor in GU. Neither in MT nor OK found matches between the MetaPhlAn and BLAST classifications.

These results show that CPF and thus, photoreactivation is usually present in the most abundant organisms of the UV_High_ microbiomes (Fig. 6). Also, SS and RA show a high number of taxa in which photoreactivation is present. That suggests that photoreactivation may act as a successful system for assuring the survival and predominance of taxa in UV-stressed environments. This fact contrasts with those less insolated samples, where natural selection doesn’t act in favor of photoreactivation promoting abundance nor diversity of taxa carrying these genes.

## 4. CONCLUSIONS

In this work, UV-B was considered for the first time as an ecological variable in a sequenced-based metagenomic study of microbial communities on a worldwide scale. Our results consistently showed concordance of high UV exposure of a given microbiome with its low microbial diversity and high CPF abundance.

We also evaluated the CPF diversity in the worldwide microbiomes and reported three novel CPF clades not identified in previous analyses. CPF was more likely present in most abundant organisms of the UV_High_ microbiomes, suggesting a significant evolutionary force for survival and dominance in highly irradiated environments. Also, cryptochrome-like genes were much more abundant in the most exposed microbiomes indicating a complementary role to standard photolyases.

Finally, metagenomics proved to be an excellent tool to reveal a clear correspondence between microbiomes, UV-exposure, and UV-B resistance mechanisms - measured in the number of gene copies. Additional methods, such as metatranscriptomics and metaproteomics, should be implemented in order to unveil the molecular dynamics of the CPF upon different light conditions.

## 5. ACKNOWLEDGEMENTS

VHA and MEF are staff researchers from the National Research Council (CONICET) in Argentina. DA is a recipient of a doctoral fellowship from CONICET. The authors have produced this manuscript despite the delays and shortages of funding execution from National Agencies in Argentina, mainly FONCyT (PICT 2013 2991) and CONICET (PIP 2015 0519, PIO-UNCA y PICT V 3825-2016), and the drastic devaluation of Argentinean currency during the period 2016-2019. This manuscript has been released as a Pre-Print at bioRxiv (Alonso-Reyes et al. 2019).

## 6. AUTHORS CONTRIBUTIONS STATEMENT

DAR designed the experiments and performed the data analysis. VA planned the research, provided training and analysis tools. DAR and VA wrote the paper. MEF and VA revised the manuscript. MEF performed HAAL sampling expeditions and provided the HAAL’s microbiomes sequences. MEF and VA obtained funding for the original project idea. All authors read and approved this manuscript.

